# Reservoir dogs: Consequences of variable contact network structures for disease spread in free-ranging dogs

**DOI:** 10.1101/2023.04.06.535810

**Authors:** Manvi Sharma, Abhijeet Kulkarni, Anjan Katna, Abi Tamim Vanak

## Abstract

The risk of disease spread is contingent on not only the biological properties of causal agents, such as viruses and bacteria, but also the socio-ecological context of the outbreak. Therefore, researchers need to incorporate the variation in animal-animal interactions in models of disease spread. Much research has focused on how ecological factors (seasonality, mating systems) can influence the variation in animal interactions, but heterogeneity in interactions may arise due to human action, and this has received little attention. We hypothesised that social-mediated differences in the landscape can have consequences for interactions for free-roaming dogs and consequently for disease spread. We used GPS collared data from 31 free-ranging dogs present along a farm-village gradient to build social contact networks to examine differences in interactions. Additionally, we simulated scenarios of rabies infection of two types – dumb and furious – on the contact network structure of village and farm dogs. We found significant differences in the node degree between village and farm contact networks, but the node betweenness was similar. Our simulations showed that infections arising in village dogs are likely to spread to that network, but infections arising in farm dogs get are less likely to spread. Interestingly, this was independent of the type of rabies infection. We discuss our results in light of control strategies for managing rabies in the tropics.

## 1. Introduction

Understanding the mechanisms underlying disease spread is important for designing control strategies (Anderson and May 1979; Rohani, Zhong, and King 2010). A mechanism-centric approach relies on quantifying the variation in animal-animal interactions and incorporating this variation in models of the spread of diseases. For example, studies show that quantifying the variation in host contact patterns can help predict the rate and the source of disease outbreaks (Keeling 1999; Chen et al. 2014). In addition to prediction, possible control strategies can be better tested for efficacy when information on how disease spreads is available. For example, culling badgers as a control strategy led to increased outbreaks in susceptible populations because of increased badger movement (Woodroffe 2006). Therefore, a mechanism-based approach to understanding disease transmission can help in explaining seemingly counterintuitive patterns of disease spread as well as in predicting disease outbreaks (Leach, Webb, and Cross 2016).

Predictive disease modelling is contingent on the availability of robust empirical data. Disease ecologists have recognised the need to explicitly incorporate animal-animal interactions while modelling the spread of diseases using network-based approaches (Silk et al. 2017; Dougherty et al. 2018; Robitaille, Webber, and Wal 2019). Social contact networks have increasingly been used to understand the seasonal variation in interactions between animals. For example, in Tasmanian devils, *Sarcophilus harrisii*, increased female-female interactions outside the mating season led to differences in the contact network structures (Hamede et al. 2009). Similarly, a study on European badgers, *Meles meles* (Weber et al. 2013) showed how stability in the social structure can ameliorate the spread of tuberculosis. Independently, purely simulation-based studies that use contact rates to test mechanisms responsible for disease persistence (Victoria J. Brookes, Dürr, and Ward 2019) or to test the efficacy of vaccination (McClure et al. 2020) have also received attention, but these studies need to be validated using empirical datasets. Studies that combine empirical and simulation approaches are critical to understand how variation in animal interactions influences disease transmission, yet not many studies combine fine-scale contact data with simulations of disease transmission.

Heterogeneity in animal interactions can arise from many factors, such as life-history traits (survival and age) (Ozella et al. 2020), seasonality in resources (Dorning and Harris 2019), or the complex mating systems that animals have evolved (Hamede et al. 2009; Ashby and Gupta 2013). In addition to these ecological causes, heterogeneity in interactions may also arise due to social causes via human presence. For example, differences in human density can lead to differences in the movement of carnivores as well as free-roaming domestic animals such as dogs (Creel et al. 2019) and have consequences for densities (Bhalla et al. 2021) that can lead to heterogeneity in interactions.

Dogs are important links in disease transmission dynamics with the wild carnivores that co-occur with them (Knobel et al. 2014; Belsare, Vanak, and Gompper 2014; Belsare and Gompper 2015). Dogs have been shown to be hosts for disease-causing viruses, such as canine distemper virus, and rabies virus that can be transmitted to wildlife that co-occur with them (Viana et al. 2015). Telemetry-based approaches are ideal for recording dog interactions as individual dogs can be tracked through the day and especially post-sunset when their activity is likely to be the highest (Maher, Ward, and Brookes 2019).

Here, we quantify the heterogeneity in interactions between free-ranging dogs that co-occur in a rural landscape in India and build an understanding of consequences for this heterogeneity for disease transmission. We tested the hypothesis that socially-mediated differences in the landscape have consequences for animal interactions in village and farm dogs. We used relocation data from 31 GPS-collared dogs to build social contact networks. Specifically, we first examined if differences in the socio-ecological landscape along a farm-village gradient can lead to different network structures in village and farm dogs. Secondly, we tested the consequences of different contact network structure on disease transmission for these two sub-populations by simulating an event of rabies infection in these different contact networks.

## 2. Methods

### 2.1 Study site

This study was conducted in and around the Shirsuphal village of Baramati Taluka of Pune District, Maharashtra in West-Central India. For further details on the study area, please see Katna et al. 2022 (Katna et al. 2022) and Tiwari et al. 2018 (Tiwari et al. 2018). The study area consists of several land-use types: irrigated and seasonal agriculture, fallows grasslands, forestry plantations, fruit orchards and other infrastructure. Commercial poultry farming, agro-pastoralism and nomadic pastoralism are the other major economic activities in the rural parts of the study area (see (Carricondo-Sanchez et al. 2019)). The study area extent (∼147 km^2^) was delineated using the outer bounds of all the telemetry locations obtained in this study (Vanak and Gompper 2007).

### 2.2 Animal Capture and Telemetry

This study spanned the period from April 2017 to September 2018. We selected a set of dogs inhabiting the main village and another set of dogs inhabiting hamlets and farm residences outside the village. We fitted GPS collars to a total of 31 dogs (which is ∼8% of the total population - Tiwari et al. 2018) of which 11 (10 males and 1 female) were classified as village dogs and 20 (19 males and 1 female) were classified as farm dogs. The dogs were restrained and captured by their respective owners. Unowned or village dogs were restrained using a net and immobilized using a combination of Ketamine and Xylazine hydrochloride (Belsare and Vanak 2013). Once restrained, each dog was fitted with a GPS collar (Africa Wildlife Tracking, South Africa) that weighed less than 5% of the animal’s body weight.

Dogs were weighed and vaccinated against rabies before release. GPS collars were set to record location data every 10 min during the breeding season and every 1 h during other periods. Location data were downloaded every 15 days using a handheld receiver provided by the collar manufacturer and uploaded to www.movebank.org.

#### Ethics approval

The ethics approval for this project was provided by the animal ethics committee of ATREE (approval number AAEC/101/2016). Permission to handle animals was obtained from dog owners or the village council.

### 2.3 Data Analysis

#### 2.3.1 Home range and movement characteristics

The animal location data were screened using the R package *move* (Kranstauber, Smolla, and Scharf 2013) for duplicate records (based on duplicate timestamps and geo-coordinates) and outliers. Site-fidelity for each individual was confirmed from the mean squared distance obtained using the R package *rhr* (Signer and Balkenhol 2015). Then, 95% home-range sizes were calculated using auto-correlated kernel density estimates (Fleming et al. 2015) using the R package *ctmm* (Calabrese, Fleming, and Gurarie 2016). We also estimated mean step lengths and daily distance travelled using the R packages *adehabitatLT* and *move*, respectively.

#### 2.3.2 Farm and village dogs

As we conducted a targeted sampling, the dogs were accordingly classified as either ‘farm’ dogs or ‘village’ dogs based on their location of capture, and information from owners/villagers. This classification was further confirmed using the centroids of the home ranges of each collared individual. The locations of the home range centroids are shown in Figure 1. The centroids of the village dog home ranges were all within the bounds of Shirsuphal village. All other dogs were classified as farm dogs.

**Figure 1.**
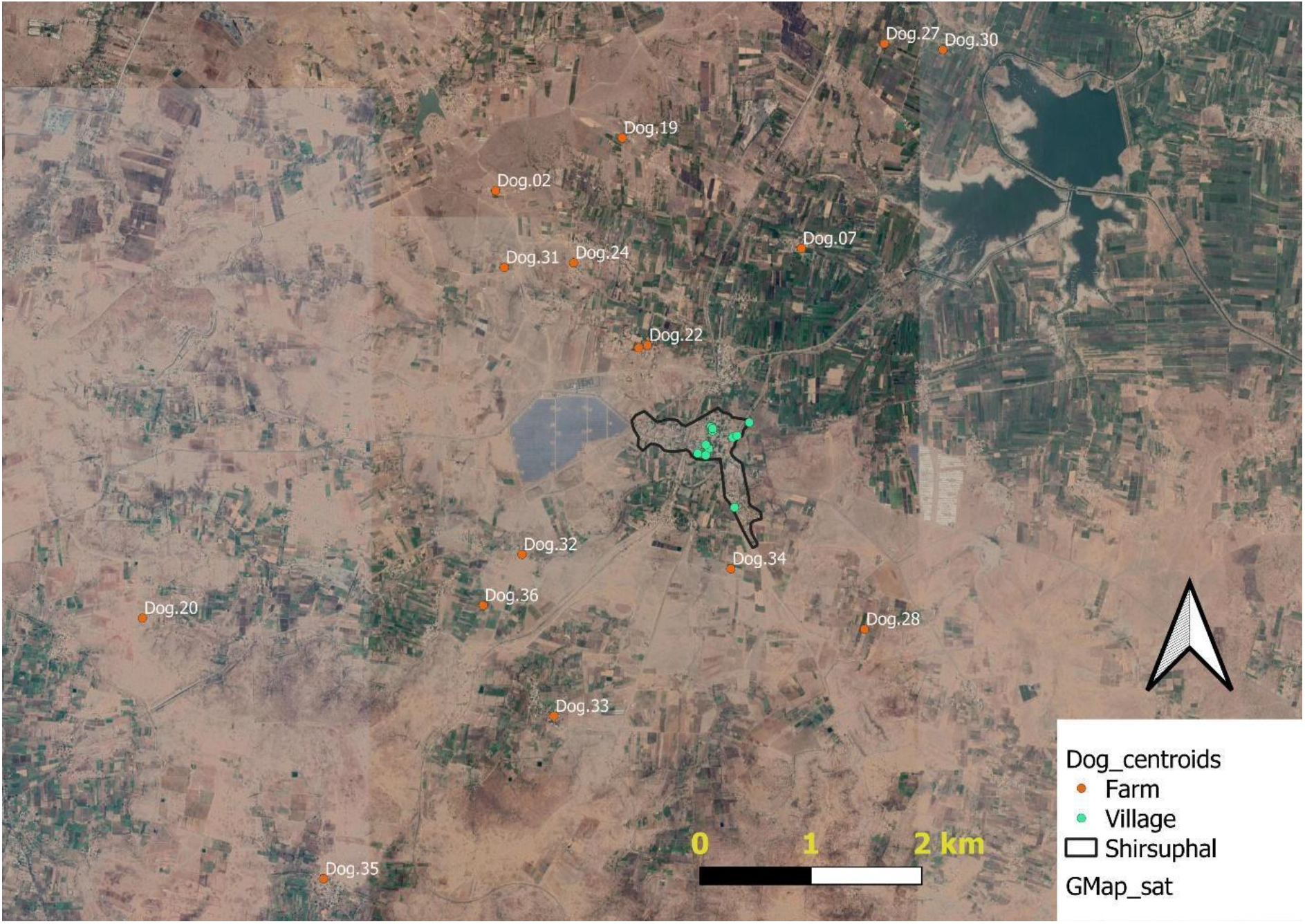
Home-range centroids of GPS-collared free-ranging dogs by category in the study site. The village boundary is marked as the black-bounding box

#### 2.3.3 Contact network analysis

We used social networks to understand the interactions between individuals in the population (Farine and Whitehead 2015; Dougherty et al. 2018). In these networks, each node represents an individual GPS-collared dog and the edges represent the interaction rate between dogs. To estimate interaction rates, studies typically use spatial overlap of estimates of home ranges, but this might not reliably translate into contact between individuals (Dougherty et al. 2018). We overcame this challenge by building proximity-based social contact networks using dog location information that clustered in both space and time. Recent advances in analytical methods account for both temporal as well as spatial proximity while building contact networks (Robitaille, Webber, and Wal 2019). We used Robitaille et al’s (2019) method of grouping relocation data. In this method, the geographic distance between animal telemetry relocations belonging to each temporal group is measured to create the networks based on proximity between animals. We created an adjacency matrix representing the connections between each dog. We calculated the average step-lengths for dogs (Figure 1 and 2) using the relocation data and used this as a threshold to define interaction between dogs (Robitaille et al 2019). In the matrix, a value of ‘1’ was assigned if the GPS location of another dog occurred within a distance corresponding to the step-length value (50m); and a value of ‘0’ was assigned if the GPS location of other dogs was farther than the step-length. In other words, the value of the interaction (edge) represented the number of other dogs within the step-length distance from a particular dog (helpfile of ‘spatsoc’ package in R, Robitaille et al. 2019). We built a global network (both village and farm dogs) and separate networks for village and farm dogs. We compared the following network properties between village and farm dogs that focussed at the individual level (node) as well the population level (network).

**Figure 2.**
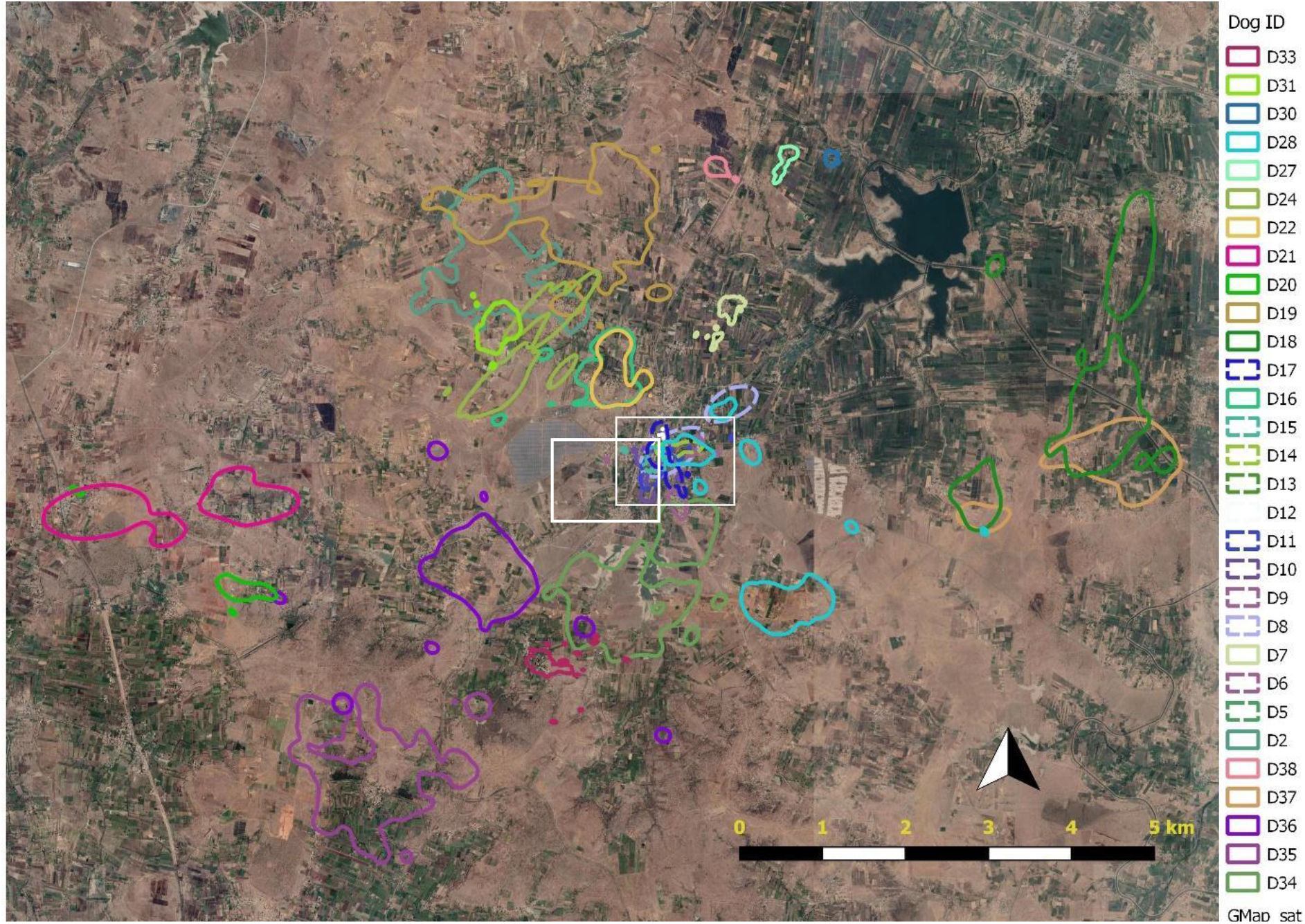
Variation in home range size of village and farm dogs. a) shows dog home ranges estimated from relocation data from collaring of 12 village dogs (inside the white box) and 19 farm dogs. b) shows a zoom-in window depicting the home range estimates from relocation data of 12 village dogs. Each colour corresponds to a unique dog that was captured, collared and released at the study site.

We calculated average network properties as described by Doughtery et al 2018 and compared it for village and farm dog networks. Node degree characterises nodes that are central to the network and represents the number of contacts per node. Node betweenness represents the frequency with which a node is on the shortest ‘path’ between pairs in the network. A high betweenness implies linking among a high number of pairs in the network and its removal, for e.g., through quarantine during an outbreak, can lead to fragmentation of the network. Edge density represents the ratio of actual links in the network and all possible links. The value ranges from 0-1, where 1 indicates that all nodes in a network are connected directly.

Given potential differences in network structures of village and farm dogs, we statistically tested if the networks show small-world properties as described by Watts and Strogatz (Watts and Strogatz 1998). Small-world networks are characterised by a high clustering coefficient and typically smaller average (shortest) path length. To test if the networks showed small-world properties, we used the standard approach of comparing clustering coefficients (cc) and average path lengths (Dube et al. 2011). We compared the clustering coefficient and average path lengths of our network with simulated draws of classical random graphs that are made by randomly creating networks with the same vertices and edges. We simulated these random networks and did permutation-based tests to compare if the clustering coefficient and average path length from our networks were significantly different from that of these networks.

All the metrics discussed above were calculated in R (R Development Core Team <u>2015</u>) using the packages *igraph* (Csardi and Nepusz 2005) and *spatsoc* (Robitaille, Webber, and Wal 2019).

#### 2.3.4 Simulations investigating the effect of dumb and furious rabies on contact rate patterns in village and farm dogs

We used a simulation based approach to investigate how the contact rate patterns in village and farm dogs get altered after an event of rabies infection. We simulated two types of rabies infection as the type of infection can result in different movement patterns in diseased dogs (Knobel et al. 2014), and that subsequently can have consequences for contact rate patterns. To simulate contact networks, we used secondary data on movement rates available from a study conducted in Zimbabwe on rabid dogs. In this study, dogs either manifested the dumb or furious form of rabies and the average distance moved per night by dogs with dumb type was four times lesser than distance moved by dogs exhibiting the furious type of rabies. The average distance moved by dogs with dumb and furious type rabies was reported to be 790m and 2885m, respectively. These data on dog movement rates were used to parametrize Brownian motion models (BMM) and generate movement trajectories for each dog from our study site. For building the BMM for rabid dogs, step-length, starting location, and turn angle values were drawn from distributions as described below. For parameterising step-lengths, we sampled from a Gaussian distribution of movement rates from Knobel et al (2013). The values for starting location were drawn from relocation data from our study site, and the values for the turn angle were drawn from a uniform distribution. Each dog infection and altered movement trajectory was simulated 50 times and the resulting (post-infection) contact networks were stored to analyse network properties. We simulated an infection with the framework that there is a fifty percent chance of developing either the dumb form or furious form of rabies. The simulated trajectories were then used to build contact networks using the same method as described in section 2.3.3. Specifically, we measured the difference in the node degree, pre- and post-infection, for each of the four scenarios described above. Similarly, we measured the differences in betweenness, pre- and post-infection, for the four scenarios. We used 95% bootstrapped confidence intervals for the interpretation of the results of the simulations.

To investigate the consequences of rabies infection on contact patterns between dogs we considered 4 scenarios: Scenario (1) Village dogs infected with dumb type virus, Scenario (2) Village dogs infected with furious type virus, Scenario (3) Farm dogs infected with dumb type rabies, and Scenario (4) Farm dogs infected with furious type rabies.

## 3. Results

### 3.1 Movement characteristics of the village and farm dogs

The mean daily distance moved by farm dogs 2.563 km (SE = 0.212) was much larger than the mean daily distance moved by village dogs 1.575 km (SE = 0.205). The mean home range estimate for farm dogs was also larger – 0.873 km^2^ (SE = 0.18) (Figure 2a) than the mean home range estimate for village dogs – 0.097 km^2^ (SE = 0.024) (Figure 2b). The mean step lengths for farm dogs were 101.89 m (SE = 0.746) and 48.232 m (SE = 0.344) for village dogs. These mean step lengths were used for building separate contact networks for village and farm dogs.

### 3.2 Variable contact networks of village and farm dogs

Overall, the network size of village and farm dogs was different. Network size, defined as the number of dogs with at least a single connection to a different dog was 100% for village dogs and 68% for farm dogs.

We estimated both individual (node) as well as population (network) level properties that are relevant to disease transmission in the village (Fig 2a) and farm dogs (Fig 2b). The histogram of the number of unique connections between nodes, showed a right-skew (skewness index = 0.82), suggesting that most nodes were connected to other nodes, while the farm dogs showed a left-skew (skewness index = -1.54), suggesting that most nodes were connected to very few other nodes.

Overall, the network structure of farm and village dogs was different at the node level as well as network level. For village dogs, the node degree was significantly higher than the node degree of the farm dog network (Figure 3a) (Permutation based statistics: difference = -15.01, p-value <0.001), suggesting that every node was well connected in the village dog network, but not in the farm dog network. For both farm and village dog networks, the node betweenness was statistically similar (Figure 3b, difference = 5.43, p-value = 0.356), suggesting the absence of any key nodes that have a relatively high number of connections with other nodes in both networks. Edge density, a property of the network, was 0.12 for the farm dog network, and 0.96 for the village dog network. A high value of edge density suggests that all nodes in a network are connected directly.

**Figure 3.**
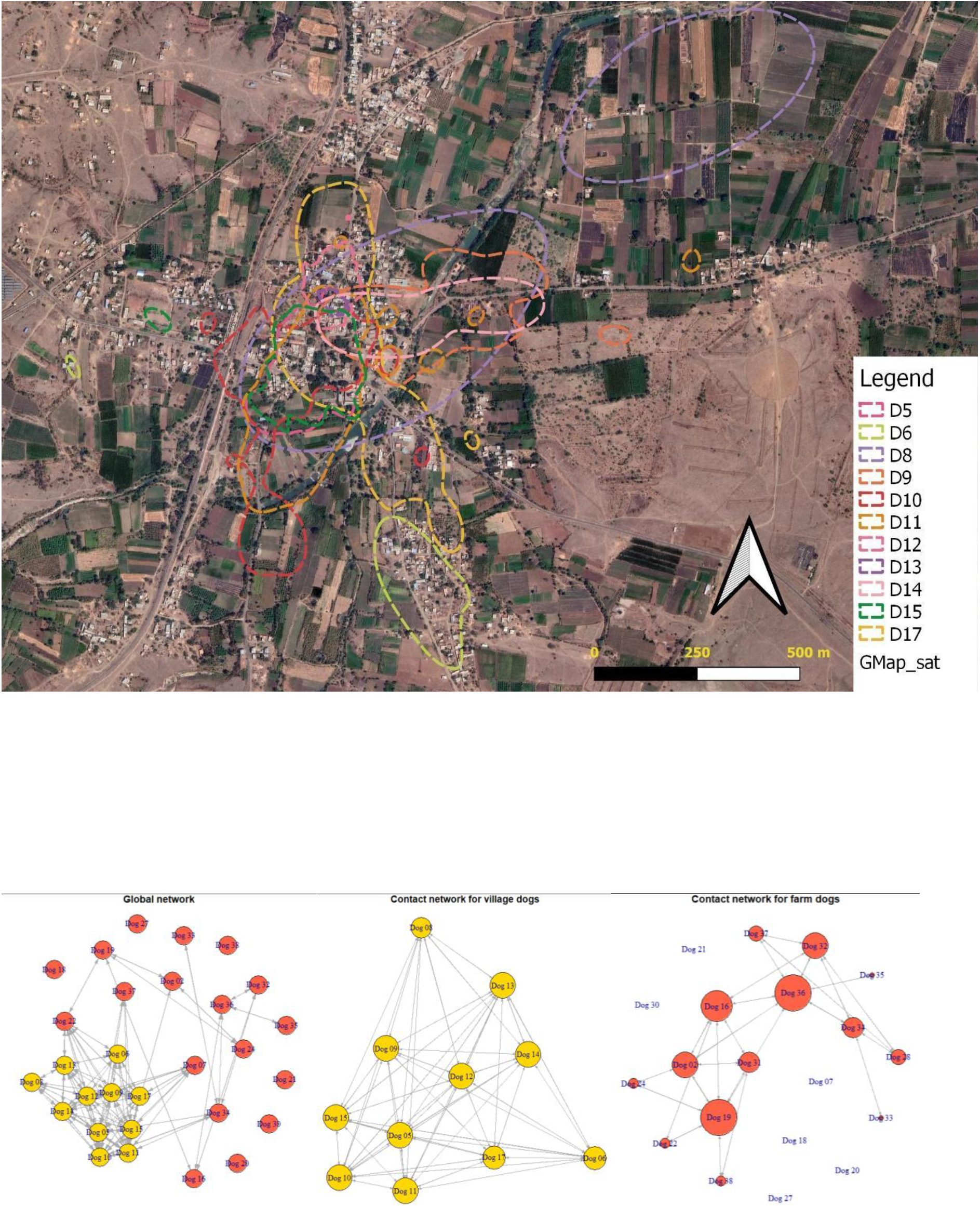
Contact networks based on spatial proximity (a) Global network with orange nodes representing farm dogs and yellow nodes representing village dogs (b) Contact network for village dogs (c) Contact network for farm dogs

**Figure 4.**
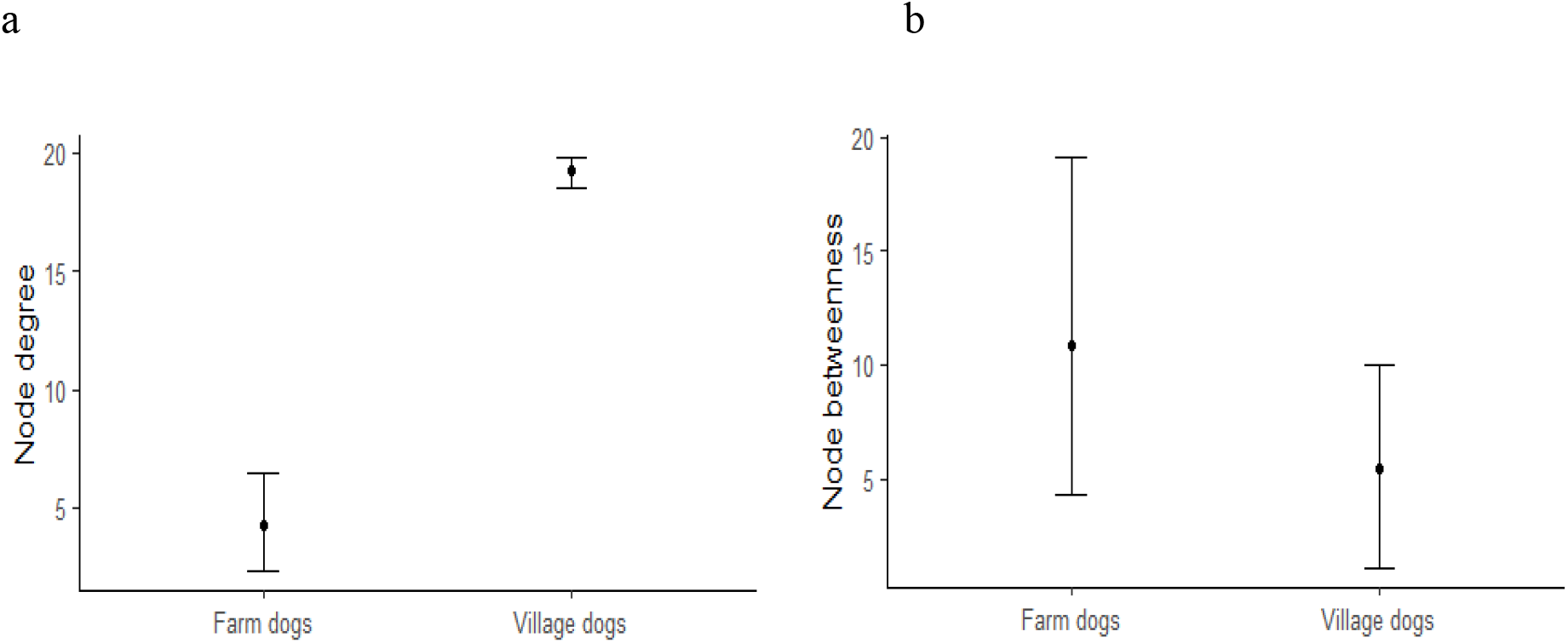
Network measures for village and farm dog networks. a) Node degree of village dog network was significantly higher than the node degree of farm dog network. b) Node betweenness for both village and farm dogs was similar. Error bars show 95% bootstrapped CIs.

**Figure 5.**
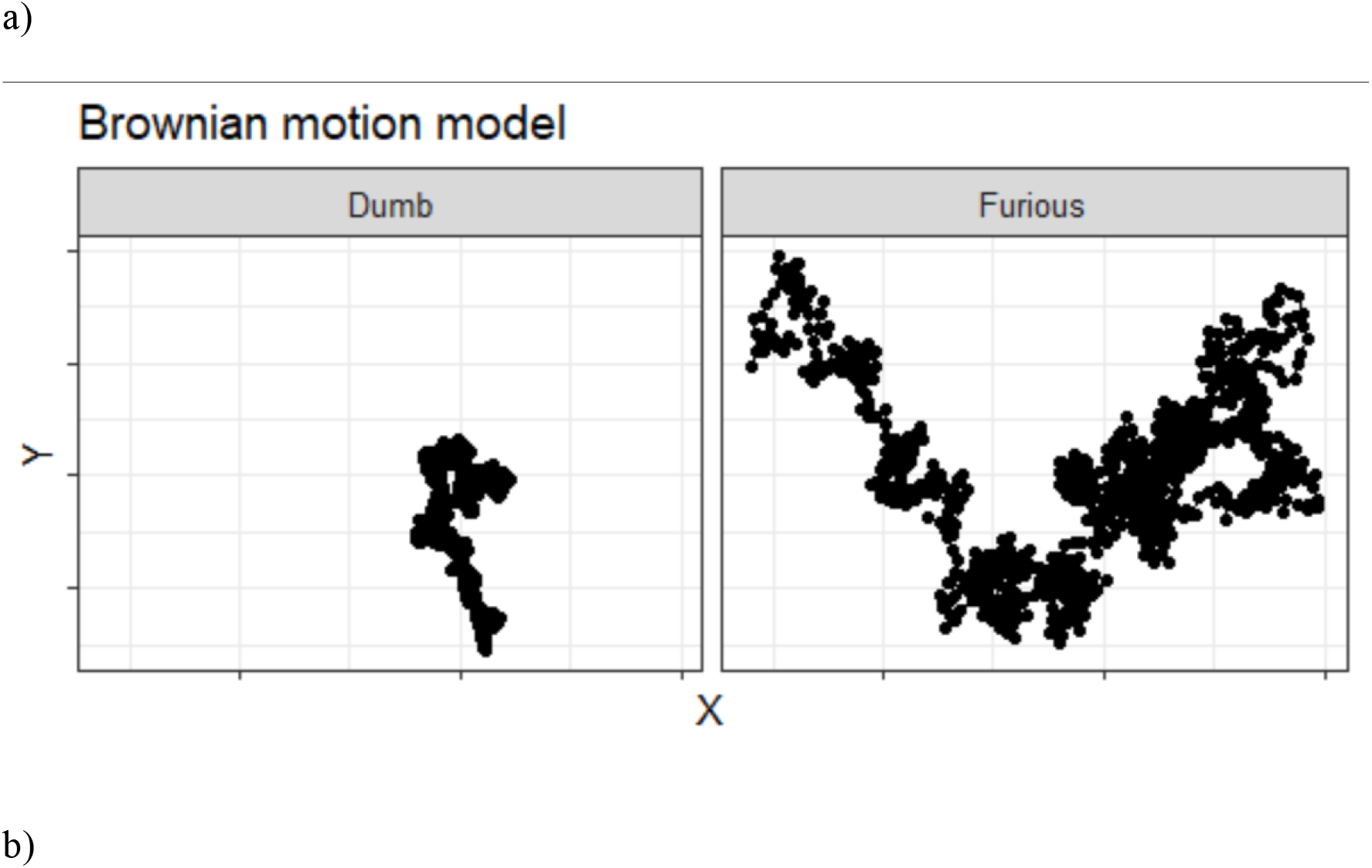
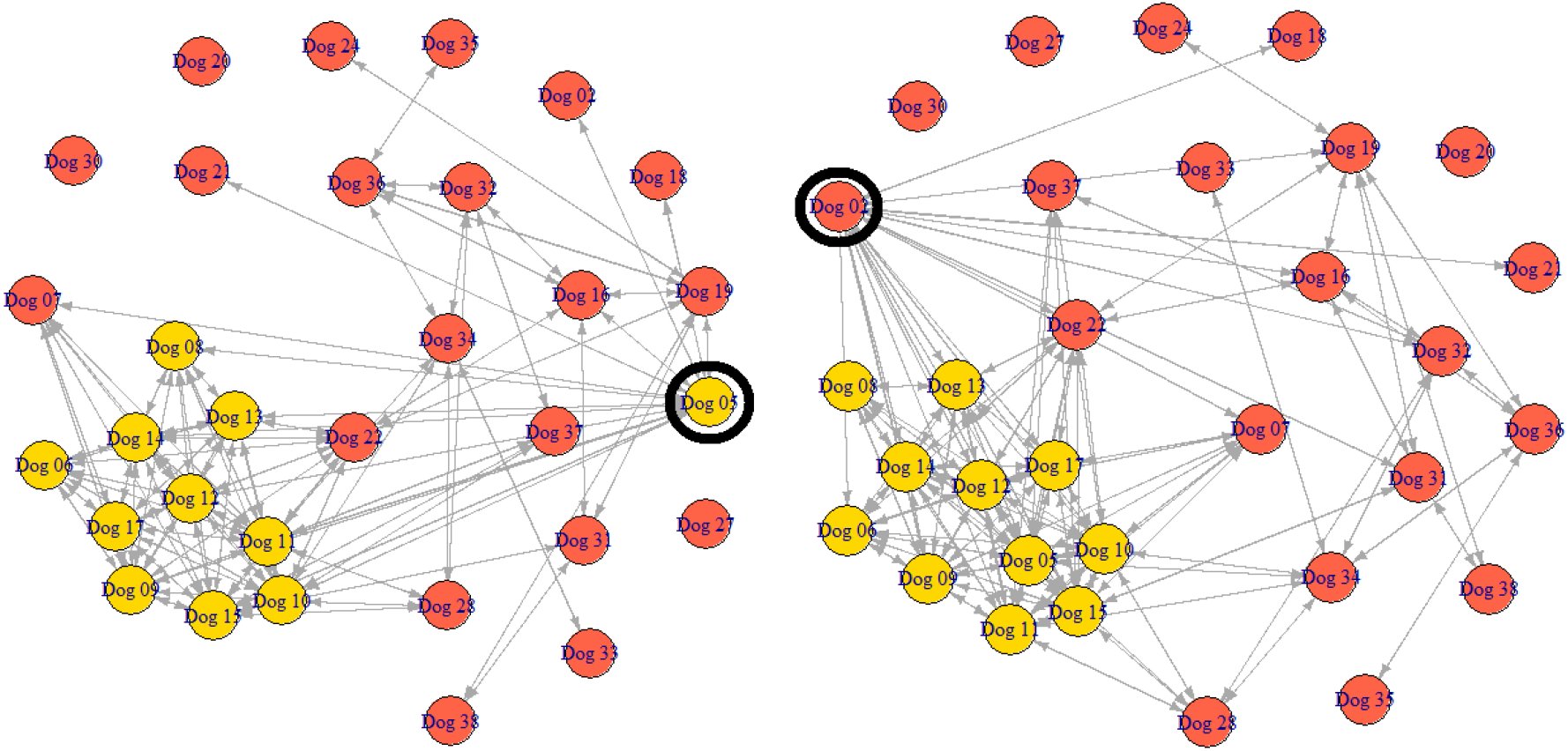
a) Simulated Brownian motion trajectories for dogs infected with dumb and furious type rabies (movement data from Knobel et al. 2014). b) Simulated contact network post-infection of a village dog (left panel) and of a farm dog (right panel). The black circles indicate an infected dog.

For the farm dog networks, we observed substantially more clustering in the observed network than expected from a random network (Permutation based test statistics (PBTS): difference in clustering = 0.38, p-value<0.001). Additionally, we found that the average of shortest paths between vertex pairs was similar to that from a random network (PBTS: difference in average path length = 0.01, p-value=0.036). The clustering coefficient of the village dog network was not significantly different from that generated from a random network with given edges and vertices (PBTS: difference in cc =0.04, p-value=0.1). The average path length was different from that of a randomly generated network (PBTS: difference in average path length = 0.03, p-value<0.001). Taken together, we find evidence for small-world behaviour in the farm dog network but the network of village dogs did not show small-world properties.

Our simulations were primarily aimed at testing if village dogs are likely to have higher rates of rabies transmission based on contact rate patterns. Secondly, we tested if the rates of transmission depended on the rabies type. Our simulations showed that in both dumb and furious rabies type of infection, the infected village dogs had a significantly larger node degree, indicative of higher contact rates, than the farm dogs (non-overlapping 95% CI in Figure 6a). However, the node betweenness, a measure of centrality, was similar between infected village and farm dogs (overlapping 95% CI, Figure 6b). We did not observe any difference in node degree or betweenness caused due to the type of rabies infection.

**Figure 6.**
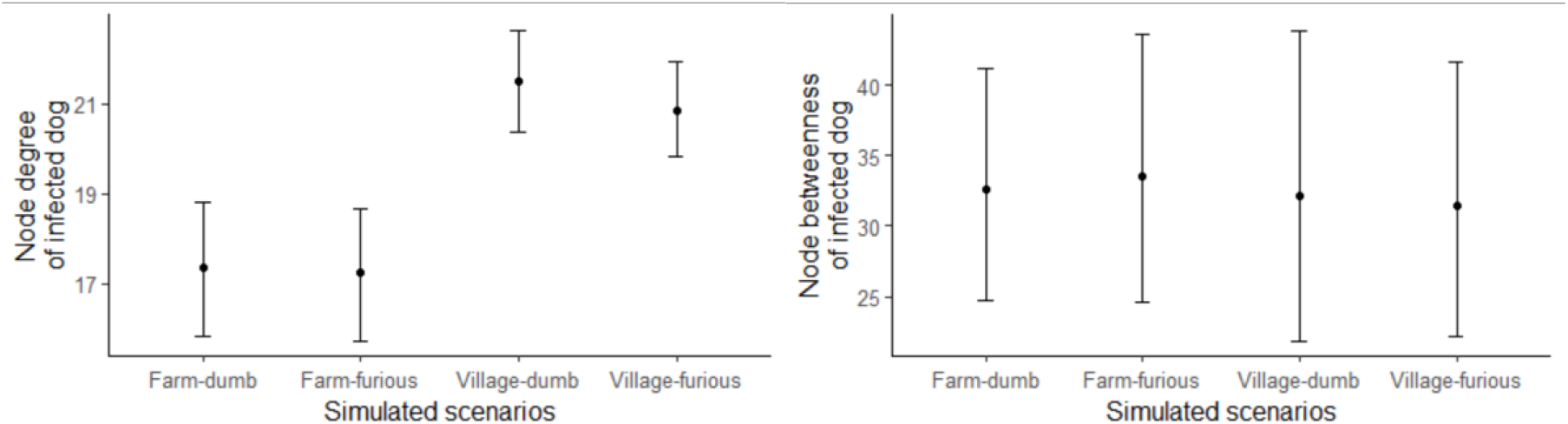
Network measures for simulated networks. a) Node degree of infected dog from the village was significantly higher than the node degree of infected dog from the farm. b) Node betweenness of both infected dogs from both village and farm was similar. Rabies type had no effect. Error bars show 95% bootstrapped CIs.

## 4. Discussion

In this study, using telemetry data and a contact network approach, we demonstrate that different dog populations can have variable network structures within a rural landscape that can have consequences for disease transmission. We find that socially mediated differences (e.g., village and farm) in the landscape can result in the dog population networks showing dramatically different properties. Through our simulations, we demonstrated how fine-scale spatio-temporal data on interacting individuals can be used to detect differences that are important for modelling contact rates in a susceptible population and can help in testing potential strategies for the control of disease spread.

India has a free-roaming dog population of about 60 million (Gompper 2014). These dogs occur in a wide variety of urban and rural matrix landscapes, often co-occurring with wild carnivore species, such as wolves, jackals and foxes. Dogs act as host reservoirs and pose a high risk of disease spill-over (Knobel et al. 2014; Belsare, Vanak, and Gompper 2014; Grover et al. 2018). Using our network-based analysis, we assessed the potential vulnerability of village dog population and farm dog population to disease spread. Farm dog networks showed small-world network properties that indicated susceptibility to the rapid spread of disease but ultimately interacted with fewer individuals. On the other hand, small-world properties were absent from village dog networks and these networks represented random mixing, suggesting high levels of interaction amongst individuals. A study conducted in Australia reported small-world properties in networks from all the communities of dogs they studied (Brookes, VanderWaal, and Ward 2020), contrary to what we find.

### Within population variation in risk of disease spread

The node degree in the village dog networks was much higher than the farm dog networks suggesting that village dogs are susceptible to frequent attacks of disease, i.e., any single dog affected is likely to rapidly affect the susceptible population. Control strategies, such as vaccination and population control, that aim at the population, and not individuals, are likely to be more successful than the quarantine of infected dogs. It is important to note here that in our study, we examined individual-node properties and only scale-free properties of networks. A critique of network-based studies is that interactions based on individuals that are sampled for building the network only represent a subset of the interactions in the population and that global network properties might be sensitive to sampling. This is less likely to affect local network properties (Dube et al. 2011). Our simulations showed that an infection that arises in a village dog is likely to spread faster than an infection that arises in a farm dog because of differentially high node degree. Surprisingly, we found that this was similarly high for both dumb and furious types of rabies despite the differences in the movement behaviour associated with the two types of rabies. Village dogs are clustered over a small area relative to the movement scale of dogs and consequently, disease transmission happens at a much smaller spatial scale. At this scale the differences that arise in movement due to rabies type are negligible. Our study confirms the prediction that it is crucial to include animal interactions accounting for the spatial scale when building models of disease transmission (Dougherty et al. 2018).

Research shows that differences in network structures in the same population across different seasons are often due to changes in mating seasons (Hamede et al. 2009). While the urban-rural gradient has been known to be an important determinant of the rate of disease spread in human health and epidemiology (Read et al. 2014), our study examines differences within rural populations networks in dogs and highlights the importance of these differences for building an understanding of disease transmission. There is a recent body of literature on the differences in the behaviour, physiology, and densities of wildlife populations between rural and urban landscapes (Batabyal, Balakrishna, and Thaker 2017; Zepeda et al. 2021). Future work needs to test hypotheses that can tease apart the factors responsible for differences in the network structures. Based on our findings, we propose that considering village and farm dogs as distinct populations, provides an appropriate scale to monitor disease spread not only because of the ecological differences but also from the point of view of disease management.

Our study shows that there is heterogeneity in dog interactions at two scales. At the broader scale, i.e., at the scale of the population of village and farm dogs, we find that interactions between networks differ, for example, differences in average node degree. These population-level properties are important in understanding the vulnerability of the population to disease spread (Spiegel et al. 2017). The second level of heterogeneity that we observed was at the level of individuals within a network. For example, in the farm dog network, we observed that certain dogs were better connected to other dogs (see Dog 19 and Dog 36 in figure 3).

This fine-scale heterogeneity between individual hosts is informative for modelling the dynamics of infection in a susceptible population, for example, estimating the R_0_ (i.e., number of secondary cases produced by a single infection in a completely susceptible population) (Dougherty et al. 2018).

Control strategies based on previous studies, that are agnostic to this heterogeneity, are likely to be inefficient and economically stressful to countries that have an already burdened economy. We suggest that control strategies that are suitable for small-world networks, such as quarantining infected dogs to fragment the network will not be effective for disease control in village dogs. Heterogeneity in other movement parameters, such as home range estimates shows that control strategies need to account for heterogeneity in movement characteristics. For e.g. village dogs have much smaller home ranges than farm dogs are likely to be more localised, but are not necessarily more accessible. On the other hand, farm dogs are more accessible (due to ownership), but require far greater effort for vaccination. Thus, our study highlights key variable parameters that need to be considered while modelling disease dynamics in dog populations. Combining these data with real-time epidemiological data can be very powerful tools for understanding disease dynamics that occur at the interface of humans and wildlife.

## References

Anderson, Roy M., and Robert M. May. 1979. “Population Biology of Infectious Diseases: Part I.” Nature 280 (5721): 361–67. https://doi.org/10.1038/280361a0.

Ashby, Ben, and Sunetra Gupta. 2013. “Sexually Transmitted Infections in Polygamous Mating Systems.” Philosophical Transactions of the Royal Society B: Biological Sciences 368 (1613): 20120048. https://doi.org/10.1098/rstb.2012.0048.

Batabyal, Anuradha, Shashank Balakrishna, and Maria Thaker. 2017. “A Multivariate Approach to Understanding Shifts in Escape Strategies of Urban Lizards.” Behavioral Ecology and Sociobiology 71 (5): 83. https://doi.org/10.1007/s00265-017-2307-3.

Belsare, A. V., A. T. Vanak, and M. E. Gompper. 2014. “Epidemiology of Viral Pathogens of Free-Ranging Dogs and Indian Foxes in a Human-Dominated Landscape in Central India.” Transboundary and Emerging Diseases 61 (1): 78–86. https://doi.org/10.1111/tbed.12265.

Belsare, Aniruddha V., and Matthew E. Gompper. 2015. “A Model-Based Approach for Investigation and Mitigation of Disease Spillover Risks to Wildlife: Dogs, Foxes and Canine Distemper in Central India.” Ecological Modelling 296 (January): 102–12. https://doi.org/10.1016/j.ecolmodel.2014.10.031.

Belsare, Aniruddha, and Abi Tamim Vanak. 2013. “Use of Xylazine Hydrochloride– Ketamine Hydrochloride for Immobilization of Indian Fox (Vulpes Bengalensis) in Field Situations.” Journal of Zoo and Wildlife Medicine 44 (3): 753–55. https://doi.org/10.1638/2012-0158R.1.

Bhalla Shireen Jagriti, Roy Kemmers, Ana Vasques, and Abi Tamim Vanak. 2021. “‘Stray Appetites’: A Socio-Ecological Analysis of Free-Ranging Dogs Living alongside Human Communities in Bangalore, India.” Urban Ecosystems 24 (6): 1245–58. https://doi.org/10.1007/s11252-021-01097-4.

Brookes, V. J., K. VanderWaal, and M. P. Ward. 2020. “The Social Networks of Free-Roaming Domestic Dogs in Island Communities in the Torres Strait, Australia.” Preventive Veterinary Medicine 181 (August): 104534. https://doi.org/10.1016/j.prevetmed.2018.09.008.

Brookes, Victoria J., Salome Dürr, and Michael P. Ward. 2019. “Rabies-Induced Behavioural Changes Are Key to Rabies Persistence in Dog Populations: Investigation Using a Network-Based Model.” PLOS Neglected Tropical Diseases 13 (9): e0007739. https://doi.org/10.1371/journal.pntd.0007739.

Calabrese, Justin M., Chris H. Fleming, and Eliezer Gurarie. 2016. “Ctmm: An r Package for Analyzing Animal Relocation Data as a Continuous-Time Stochastic Process.” Methods in Ecology and Evolution 7 (9): 1124–32. https://doi.org/10.1111/2041-210X.12559.

Carricondo-Sanchez, David, Morten Odden, Abhijeet Kulkarni, and Abi Tamim Vanak. 2019. “Scale-Dependent Strategies for Coexistence of Mesocarnivores in Human-Dominated Landscapes.” Biotropica 51 (5): 781–91. https://doi.org/10.1111/btp.12705.

Chen, Shi, Brad J. White, Michael W. Sanderson, David E. Amrine, Amiyaal Ilany, and Cristina Lanzas. 2014. “Highly Dynamic Animal Contact Network and Implications on Disease Transmission.” Scientific Reports 4 (1): 4472. https://doi.org/10.1038/srep04472.

Creel, Scott, Göran Spong, Matthew Becker, Chuma Simukonda, Anita Norman, Bastian Schiffthaler, and Clive Chifunte. 2019. “Carnivores, Competition and Genetic Connectivity in the Anthropocene.” Scientific Reports 9 (1): 16339. https://doi.org/10.1038/s41598-019-52904-0.

Csardi, Gabor, and Tamas Nepusz. 2005. “The Igraph Software Package for Complex Network Research.” InterJournal Complex Systems (November): 1695.

Dorning, Jo, and Stephen Harris. 2019. “Individual and Seasonal Variation in Contact Rate, Connectivity and Centrality in Red Fox (Vulpes Vulpes) Social Groups.” Scientific Reports 9 (1): 20095. https://doi.org/10.1038/s41598-019-56713-3.

Dougherty, Eric R., Dana P. Seidel, Colin J. Carlson, Orr Spiegel, and Wayne M. Getz. 2018. “Going through the Motions: Incorporating Movement Analyses into Disease Research.” Edited by Kevin Lafferty. Ecology Letters 21 (4): 588–604. https://doi.org/10.1111/ele.12917.

Dube, C., C.S. Ribble, D. Kelton, and B. McNAB. 2011. “Introduction to Network Analysis and Its Implications for Animal Disease Modelling: -EN--FR-Introduction à l’analyse Des Réseaux et à Ses Conséquences Pour La Modélisation de La Santé Animale -ES-Introducción al Análisis de Redes y Sus Consecuencias Para La Elaboración de Modelos de Enfermedades Animales.” Revue Scientifique et Technique de l’OIE 30 (2): 425–36. https://doi.org/10.20506/rst.30.2.2043.

Farine, Damien R., and Hal Whitehead. 2015. “Constructing, Conducting and Interpreting Animal Social Network Analysis.” Journal of Animal Ecology 84 (5): 1144–63. https://doi.org/10.1111/1365-2656.12418.

Fleming, C. H., W. F. Fagan, T. Mueller, K. A. Olson, P. Leimgruber, and J. M. Calabrese. 2015. “Rigorous Home Range Estimation with Movement Data: A New Autocorrelated Kernel Density Estimator.” Ecology 96 (5): 1182–88. https://doi.org/10.1890/14-2010.1.

Grover, Michael, Paul R. Bessell, Anne Conan, Pim Polak, Claude T. Sabeta, Bjorn Reininghaus, and Darryn L. Knobel. 2018. “Spatiotemporal Epidemiology of Rabies at an Interface between Domestic Dogs and Wildlife in South Africa.” Scientific Reports 8 (1): 10864. https://doi.org/10.1038/s41598-018-29045-x.

Hamede, Rodrigo K., Jim Bashford, Hamish McCallum, and Menna Jones. 2009. “Contact Networks in a Wild Tasmanian Devil (Sarcophilus Harrisii) Population: Using Social Network Analysis to Reveal Seasonal Variability in Social Behaviour and Its Implications for Transmission of Devil Facial Tumour Disease.” Ecology Letters 12 (11): 1147–57. https://doi.org/10.1111/j.1461-0248.2009.01370.x.

Hudson, Emily G., Victoria J. Brookes, Salome Dürr, and Michael P. Ward. 2019. “Targeted Pre-Emptive Rabies Vaccination Strategies in a Susceptible Domestic Dog Population with Heterogeneous Roaming Patterns.” Preventive Veterinary Medicine 172 (November): 104774. https://doi.org/10.1016/j.prevetmed.2019.104774.

Katna, A., A. Kulkarni, M. Thaker, and A. T. Vanak. 2022. “Habitat Specificity Drives Differences in Space-Use Patterns of Multiple Mesocarnivores in an Agroecosystem.” Journal of Zoology 316 (2): 92–103.

Keeling, Matthew J. 1999. “The Effects of Local Spatial Structure on Epidemiological Invasions.” Proceedings of the Royal Society of London. Series B: Biological Sciences 266 (1421): 859–67.

Knobel, Darryn L., James RA Butler, Tiziana Lembo, Rob Critchlow, and Matthew E. Gompper. 2014. Free-Ranging Dogs and Wildlife Conservation. OUP Oxford.

Kranstauber, Bart, Marco Smolla, and A. K. Scharf. 2013. “Move: Visualizing and Analyzing Animal Track Data.” R Package Version 1 (360): r365.

Leach, Clint, Colleen T. Webb, and Paul C. Cross. 2016. “When Environmentally Persistent Pathogens Transform Good Habitat into Ecological Traps.” Royal Society Open Science. https://doi.org/10.1098/rsos.160051.

Maher, Elizabeth K., Michael P. Ward, and Victoria J. Brookes. 2019. “Investigation of the Temporal Roaming Behaviour of Free-Roaming Domestic Dogs in Indigenous Communities in Northern Australia to Inform Rabies Incursion Preparedness.” Scientific Reports 9 (1): 14893. https://doi.org/10.1038/s41598-019-51447-8.

Ozella, Laura, Joss Langford, Laetitia Gauvin, Emily Price, Ciro Cattuto, and Darren P. Croft. 2020. “The Effect of Age, Environment and Management on Social Contact Patterns in Sheep.” Applied Animal Behaviour Science 225 (April): 104964. https://doi.org/10.1016/j.applanim.2020.104964.

Read, Jonathan M., Justin Lessler, Steven Riley, Shuying Wang, Li Jiu Tan, Kin On Kwok, Yi Guan, Chao Qiang Jiang, and Derek A. T. Cummings. 2014. “Social Mixing Patterns in Rural and Urban Areas of Southern China.” Proceedings of the Royal Society B: Biological Sciences 281 (1785): 20140268. https://doi.org/10.1098/rspb.2014.0268.

Robitaille, Alec L., Quinn M. R. Webber, and Eric Vander Wal. 2019. “Conducting Social Network Analysis with Animal Telemetry Data: Applications and Methods Using Spatsoc.” Methods in Ecology and Evolution 10 (8): 1203–11. https://doi.org/10.1111/2041-210X.13215.

Rohani, Pejman, Xue Zhong, and Aaron A. King. 2010. “Contact Network Structure Explains the Changing Epidemiology of Pertussis.” Science 330 (6006): 982–85.

Signer, Johannes, and Niko Balkenhol. 2015. “Reproducible Home Ranges (Rhr): A New, User-Friendly R Package for Analyses of Wildlife Telemetry Data.” Wildlife Society Bulletin 39 (2): 358–63. https://doi.org/10.1002/wsb.539.

Silk, Matthew J., Darren P. Croft, Richard J. Delahay, David J. Hodgson, Mike Boots, Nicola Weber, and Robbie A. McDonald. 2017. “Using Social Network Measures in Wildlife Disease Ecology, Epidemiology, and Management.” BioScience 67 (3): 245–57. https://doi.org/10.1093/biosci/biw175.

Sparkes, Jessica, Guy Ballard, Peter J. S. Fleming, Remy van de Ven, and Gerhard Körtner. 2016. “Contact Rates of Wild-Living and Domestic Dog Populations in Australia: A New Approach.” Oecologia 182 (4): 1007–18. https://doi.org/10.1007/s00442-016-3720-4.

Tiwari, Harish Kumar, Abi Tamim Vanak, Mark O’Dea, and Ian Duncan Robertson. 2018. “Knowledge, Attitudes and Practices towards Dog-Bite Related Rabies in Para-Medical Staff at Rural Primary Health Centres in Baramati, Western India.” PloS One 13 (11): e0207025.

Vanak, Abi Tamim, and Matthew E. Gompper. 2007. “Effectiveness of Non-Invasive Techniques for Surveying Activity and Habitat Use of the Indian Fox Vulpes Bengalensis in Southern India.” Wildlife Biology 13 (2): 219–24.

Viana, Mafalda, Sarah Cleaveland, Jason Matthiopoulos, Jo Halliday, Craig Packer, Meggan E. Craft, Katie Hampson, et al. 2015. “Dynamics of a Morbillivirus at the Domestic– Wildlife Interface: Canine Distemper Virus in Domestic Dogs and Lions.” Proceedings of the National Academy of Sciences 112 (5): 1464–69. https://doi.org/10.1073/pnas.1411623112.

Watts, Duncan J., and Steven H. Strogatz. 1998. “Collective Dynamics of ‘Small-World’ Networks.” Nature 393 (6684): 440–42. https://doi.org/10.1038/30918.

Weber, Nicola, Stephen P. Carter, Sasha R. X. Dall, Richard J. Delahay, Jennifer L. McDonald, Stuart Bearhop, and Robbie A. McDonald. 2013. “Badger Social Networks Correlate with Tuberculosis Infection.” Current Biology 23 (20): R915–16. https://doi.org/10.1016/j.cub.2013.09.011.

Woodroffe, r. 2006. “Effects of Culling on Badger Meles Meles Spatial Organization: Implications for the Control of Bovine Tuberculosis.” Journal of Applied Ecology. https://besjournals.onlinelibrary.wiley.com/doi/full/10.1111/j.1365-2664.2005.01144.x.

Zepeda, Emily, Eric Payne, Ashley Wurth, Andrew Sih, and Stanley Gehrt. 2021. “Early Life Experience Influences Dispersal in Coyotes (Canis Latrans).” Behavioral Ecology 32 (4): 728–37. https://doi.org/10.1093/beheco/arab027.

